# Network- and Enrichment-based Inference of Phenotypes and Targets from large-scale Disease Maps

**DOI:** 10.1101/2021.09.13.460023

**Authors:** Matti Hoch, Suchi Smita, Konstantin Cesnulevicius, David Lescheid, Myron Schultz, Olaf Wolkenhauer, Shailendra Gupta

## Abstract

Disease maps have emerged as computational knowledge bases for exploring and modeling disease-specific molecular processes. By capturing molecular interactions, disease-associated processes, and phenotypes in standardized representations, disease maps provide a platform for applying bioinformatics and systems biology approaches. Applications range from simple map exploration to algorithm-driven target discovery and network perturbation. The web-based MINERVA environment for disease maps provides a platform to develop tools not only for mapping experimental data but also to identify, analyze and simulate disease-specific regulatory networks. We have developed a MINERVA plugin suite based on network topology and enrichment analyses that facilitate multi-omics data integration and enable *in silico* perturbation experiments on disease maps. We demonstrate workflows by analyzing two RNA-seq datasets on the Atlas of Inflammation Resolution (AIR). Our approach improves usability and increases the functionality of disease maps by providing easy access to available data and integration of self-generated data. It supports efficient and intuitive analysis of omics data, with a focus on disease maps.

## INTRODUCTION

### Background

Molecular and cell biology has amassed a tremendous amount of information on molecular interactions related to disease development, progression, and treatment. Clinical scientists and biomedical researchers have access to any chosen disease phenotype, process, or molecule through databases built on scientific literature and experimental data. However, searching publications and databases for molecules of interest, identifying regulatory mechanisms and potential drug targets is, in most practical cases, a long-term research project rather than a quick task that supports the diagnosis and treatment of patients.

### The Disease Map Approach

#### Disease maps

are developed to support the disease-oriented exploration of state-of-the-art knowledge. Community-built disease maps are comprehensive and accessible resources that collect validated knowledge about a disease, its molecules, phenotypes, and processes (Mazein *et al*, 2018b; Ostaszewski et al, 2019). Encoding this knowledge in a standardized format enables established analytical tools to extract information from the complex interactions or perform *in silico* experiments on integrated experimental data (Figure 1). Examples of published disease maps include the Parkinson’s Disease Map (Fujita et al, 2014), the Rheumatoid Arthritis Map (Singh et al, 2018), the AsthmaMap (Mazein et al, 2018a), or the Atherosclerosis Map (Parton et al, 2019).

**Figure 1:**
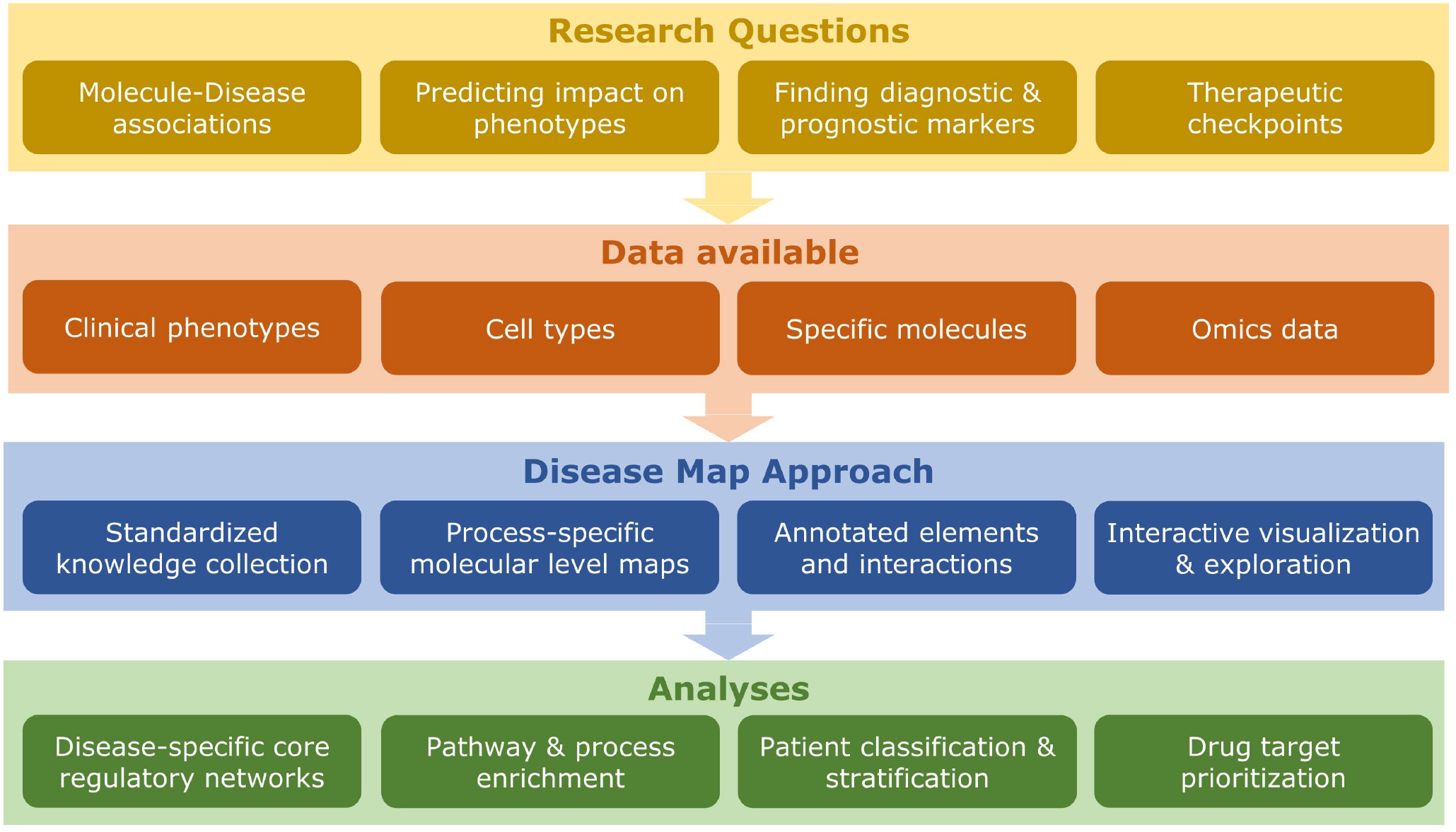
Overview of the Disease Map Approach to help to address disease-specific research questions, as it is implemented with the plugin suite.

Systems biology standards for encoding contextual information, such as Systems Biology Markup Language (SBML), encoding visual information, such as Systems Biology Graphical Notation (SBGN), or encoding both information simultaneously, such as CellDesigner-SBML, can be used to organize molecular interactions into submaps and layers (Keating *et al*, 2020; Kitano *et al*, 2005; Funahashi *et al*, 2007). A **submap** describes the molecular interactions that regulate related biological processes or clinically observable signs and symptoms, usually represented as SBGN **phenotype** elements. Elements of these maps are linked to public databases and organized into different layers that aid in the visualization and exploration of disease maps. Figure 2 gives an example of a submap from the “Atlas of Inflammation Resolution” (AIR) (Serhan *et al*, 2020).

**Figure 2:**
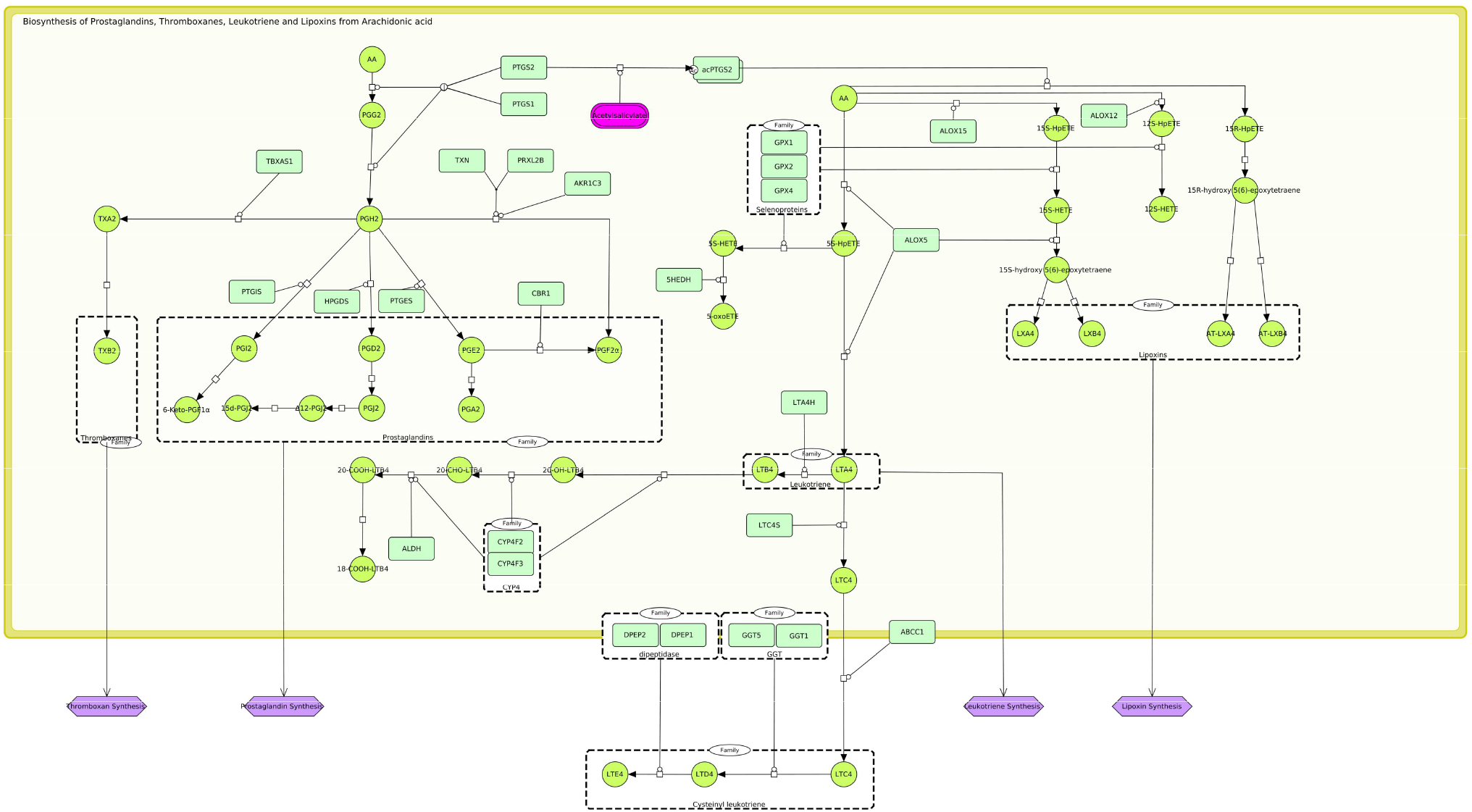
SBGN representation of the “Biosynthesis of PIM and SPM from AA” in the “Atlas of Inflammation Resolution” (AIR). Molecular interactions represented in the SBML process description format that are involved in the regulation of various phenotypes (purple) such as “thromboxane synthesis” or “prostaglandin synthesis.”

The curation of submaps is a manual process that aggregates experimentally validated evidence from the literature and provides a rich annotation of interactions with links to various databases. To make such maps publicly accessible and interactive to the community, **MINERVA** was developed as a web-based platform for curating and interactively visualizing maps that support community-driven projects (Gawron et al, 2016). Due to automated annotation with multiple databases and extensive exploration tools, many of the currently published disease maps, including the AIR, are hosted on MINERVA. In the AIR, the submaps are additionally enriched with protein-protein interactions (PPI) and regulatory information, including transcription factors (TF), microRNA (miRNA), or long non-coding RNA (lncRNA) interactions. The entirety of molecular interactions, the “bottom layer” of the disease map, combining information from submaps and regulatory interactions, we refer to as the **molecular interaction map (MIM)**. The MIM encodes information about molecules and their interactions in pathways, networks, and their relationship to disease phenotypes. Even for a narrowly defined context, most disease maps will include large numbers of interactions. By mapping omics data to corresponding elements in the MIM, experimental measurements such as changes in gene expression, metabolite concentrations, or genetic mutations can be integrated into the disease map. The integration of differential omics data is highly interesting for users to identify regulatory elements and processes or predict the consequences of changes induced in the MIM. A disease map provides a platform for analyzing experimental data and inferring interaction paths from maps by focusing on chosen disease phenotypes.

### Research gap

To date, intuitive analysis tools for disease maps have been limited and typically require implementing external tools into the workflow. Consequently, data must be exported, transformed, and imported, which requires knowledge of programming languages and limits the usability of disease maps for non-bioinformaticians. Nonetheless, many publicly available tools enable topological analysis, target prediction, motif identification, and more based on standard molecular interaction maps. The recently published CasQ converts SBML maps curated in process description (PD) format into qual-SBML, a file format for executable Boolean models (Aghamiri *et al*, 2020). The generated models can then be imported into stand-alone or web-based tools such as CellCollective for further analyses (Helikar *et al*, 2012). Many common tools for data analysis are based on enrichment approaches, most commonly the “Gene Set Enrichment Analysis” (GSEA). The GSEA statistically evaluates the probability that a user-supplied list of genes is overrepresented in defined gene sets related to a clinical or biological term (Subramanian *et al*, 2005). Several commonly used analysis tools have integrated the enrichment approach, such as Enrichr (Chen *et al*, 2013), DAVID (Jiao *et al*, 2012), or ClueGO (Bindea *et al*, 2009). The Enrichr web platform provides a simple user interface and uses many public databases for its gene sets, including pathway resources such as KEGG or WikiPathways (Chen *et al*, 2013; Martens *et al*, 2021; Kanehisa, 2000). However, the results of established enrichment analyses are limited in their interpretability. They provide a statistical evaluation of whether a pathway may be overrepresented in the data, but less information about (i) the type of regulation (up- or down-regulation), (ii) the relationships between genes and terms, (iii) the range of fold changes, or (iv) the importance of each gene (its weighting) in the set. In the years following the establishment of GSEA, numerous adaptations of enrichment approaches have addressed these limitations to broaden their scope for specific purposes. Modern methods distinguish between up- and down-regulated genes (Hong *et al*, 2014; Warden *et al*, 2013) or integrate network topology information into their algorithms (Zito *et al*, 2021). In 2014, QIAGEN published the “Ingenuity Pathway Analysis” (IPA) software that provides a range of network-based solutions to infer knowledge from molecular data (Krämer *et al*, 2014). IPA is similar to the disease maps approach because it visualizes molecular pathways and provides data integration and analysis tools. Still, it has been designed for commercial use, lacking the expandability and flexibility for community-driven projects. However, these tools are usually limited to a specific case, and it can be complicated to find the most appropriate one for the desired purpose. These limitations affect the integration of data, forcing the user to import the data manually and then export the results to the disease map for visualization. Thus, there is a need for tools that enable *in silico* experiments, data integration, and data visualization directly on disease maps through intuitive and straightforward user interface elements. Given the complex nature of disease maps and the wide range of applications that these tools would need to encompass, there are many challenges to their development, some of which we summarized in Box 1:

#### Box 1: Challenges for the development of tools for knowledge inference from large-scale disease maps.

**Size:** To identify new regulators and to accurately integrate-omics datasets, disease maps contain a vast collection of molecular interactions. Established mechanistic modeling approaches, for example ordinary differentially equations (ODEs), are not well suitable for large-scale maps.

**Diversity:** Disease maps should represent the molecular pathways of the phenotypes as accurately as possible. That includes accounting for various types of molecules like metabolites, proteins, or noncoding RNAs (ncRNAs). Additionally, their interaction can be remarkably diverse, including allosteric regulations, catalysis, complex formation, transport processes, or degradation. Moreover, the types of data supplied by the users are just as multifarious as the curated molecular interactions in the map. Omics data such as genomics, transcriptomics, proteomics, or metabolomics have different data formats and require different analysis approaches.

**Userbase:** Disease maps are created to help wet lab researchers and clinicians to understand the disease pathophysiology. The inference of information should not depend on the installation of tools or the knowledge of programming languages. Ideally, tools should be fast and easily executable on many devices, preferably through a web browser with an intuitive user interface. Therefore, they must be designed for low processing power and should require as little specifications by the user as possible.

**Security:** Many datasets, especially from clinicians, fall under extremely strict data security laws. If possible, tools should prevent any upload or storage of information.

**Standards:** Many modeling approaches are applicable to a specific representation of SBML only. Whereas Boolean models require logical rules, i.e., maps in qual-SBML format, kinetic models are bound to mechanistic representations of interactions. Therefore, the curation of the map influences the analytic possibilities.

### Outcomes

To address these challenges, we developed enrichment- and network-based approaches that, through the combination of topology and data-integration methods, facilitate deriving information from disease maps. Since MINERVA supports customized plugins that can interact with the displayed submaps, it provides an excellent framework for community-driven and application-focused projects. By integrating our approaches in a multifunctional, interactive MINERVA plugin suite within the AIR, we help users answer their research questions on the map itself and visualize results in colored overlays of map elements. To demonstrate applications of the tools, we derived regulated phenotypes from a bulk RNA-seq dataset of a murine colitis model (Czarnewski et al, 2019). Additionally, we applied the target prediction algorithm to an RNA-seq dataset of IFNα-stimulated B cells and identified well-known downstream effectors of IFNα as targets (Mostafavi et al, 2016). Both case studies demonstrated the successful identification of known key targets and regulated processes.

More information and detailed user guides for the plugins are available at: https://air.bio.informatik.uni-rostock.de/plugins

## RESULTS

### A plugin suite for disease map knowledge inference

We present a suite of MINERVA plugins, initially developed for the Atlas Inflammation Resolution (AIR) but adaptable for other disease maps. The plugins can be accessed directly from the AIR (https://air.elixir-luxembourg.org/) and are thus easily accessible from any web browser (Figure **3**A).

**Figure 3:**
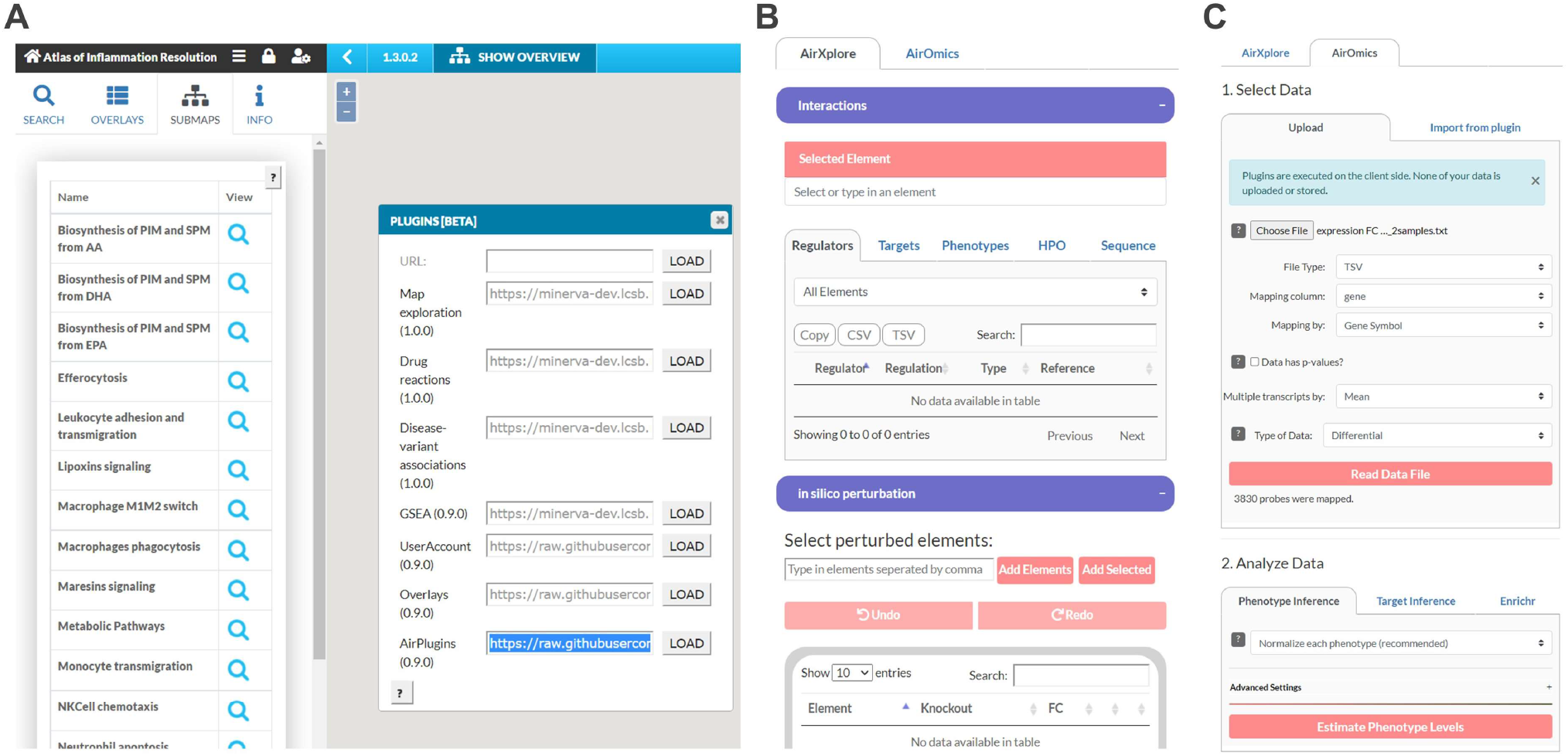
User Interface (UI) of the AirPlugins. Screenshots of how to initialize the plugins on the AIR (A) and the initial UIs of the AirXplore (B) and AirOmics (C) plugins.

The two central components of the plugin suite are the Xplore and Omics plugins, both of which include multiple tools in an intuitive user interface that enable user interactions, *in silico* experimentation, and data analysis on disease maps. We adapted and combined established enrichment algorithms with causal information from large-scale disease map models to develop sophisticated research tools: The Xplore plugin (Figure **3**B) enables data-independent in silico perturbation experiments and exploration of information stored in the MIM, whereas the Omics plugin (Figure **3**C) enables statistically evaluated inference of regulated processes and disease targets from context-specific omics data. Table 1 gives an overview of the main features of both plugins.

**Table 1:**
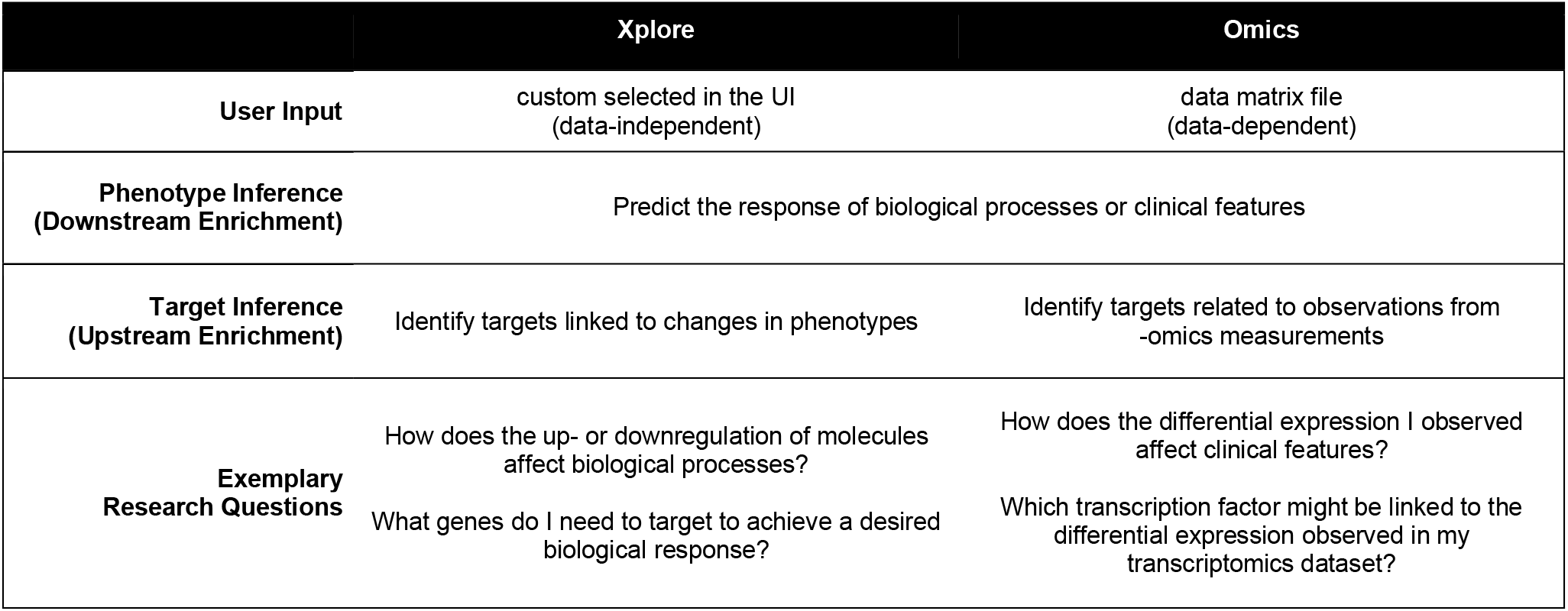
Comparison of the Xplore and Omics plugins for data-independent and data-dependent disease map analysis.

### Two-dimensional network-based enrichment analysis

Gene enrichment analyses evaluate whether a number of differentially expressed genes (DEGs), commonly referred to as the **gene list**, are statistically overrepresented in an *a priori* defined set of genes associated with a biological or clinical **term**. We have extended the definition of the gene list as input to a set of arbitrary molecules with quantitative (level-based) or qualitative (activity-based) changes, for which we introduce the notion of **differentially changed elements (DCE)**. DCEs are elements characterized by a significant *log2* fold change value (FC), either derived from transcriptomics, proteomics, or metabolomics experiments (**data-dependent** DCEs) or simply assumed by the user (**data-independent**, *in silico* simulated DCEs). DCEs can also be phenotypes referring to increased (positive value) or decreased (negative value) activities of measurable biological processes or clinical features. Additionally, we redefined the term to be enriched as any element in the MIM that is either regulated by the DCEs or is self-regulated, resulting in two main types of analyses: downstream and upstream enrichment, as summarized in Figure 4.

**Figure 4:**
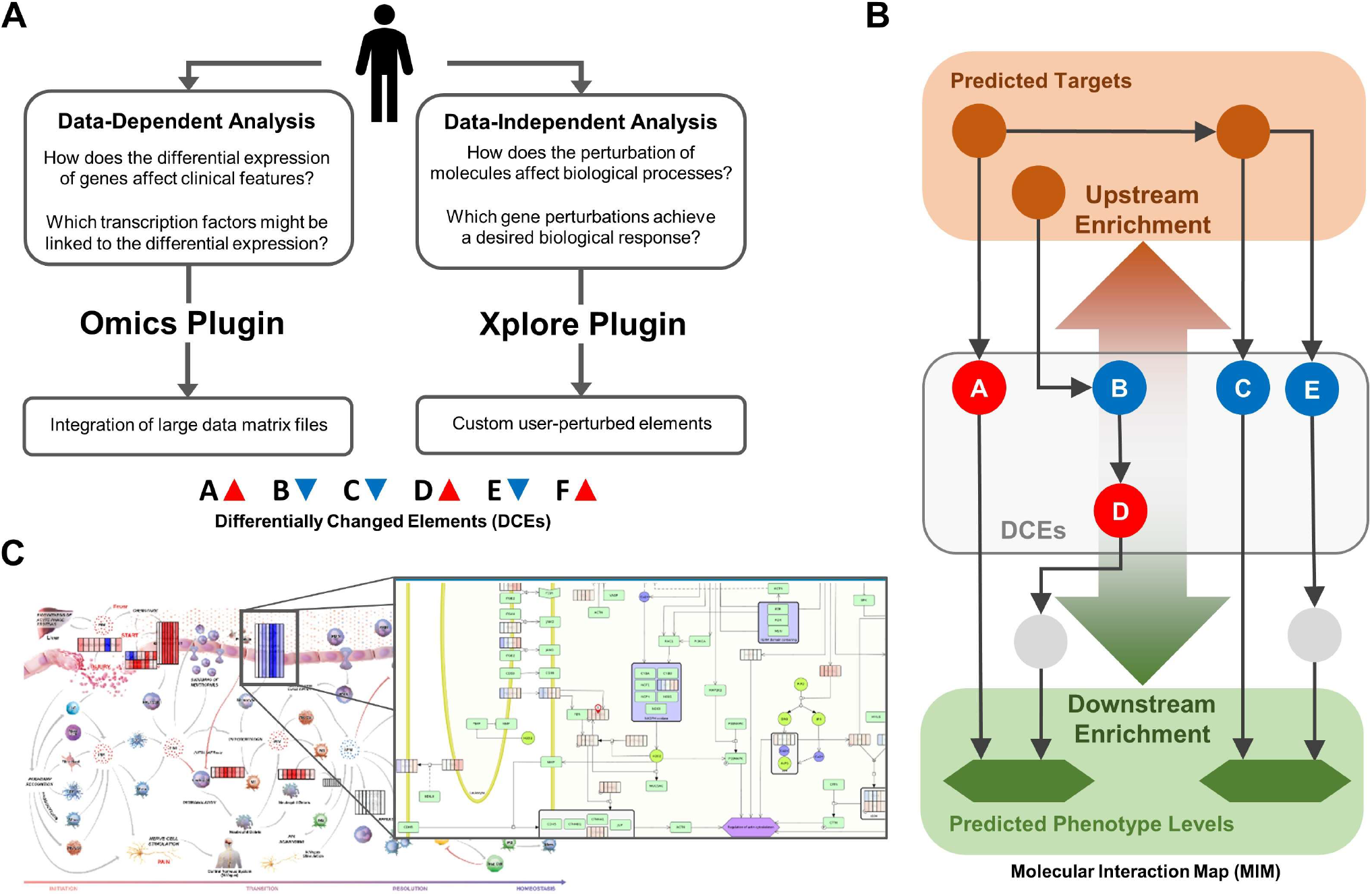
Derivation of targets and regulated phenotypes from disease map integrated-omics data through back- and forward tracing of causal interactions from DCEs. (A) Dependent on the research questions, users can either upload data files to the Omics plugin or manually perturb specific elements *in silico* using the Xplore plugin. (B) From the set of differentially changed elements (DCEs), the network-based enrichment can statistically evaluate their impact on phenotypes or their regulation from potential targets in the MIM. (C) Results are visualized as colored overlays on the submaps to enable intuitive interpretations.

We applied network topology algorithms to express the relationship between each pair of elements in the whole MIM as a numerical value called the **influence score**. These scores provide a second weighting factor besides the fold change value, which describes the impact of the DCEs on the enriched term. Influence scores depend on the context and origin of the data. Transcriptional influence describes the effect of a MIM element on the transcription of a particular gene in transcriptomics data analyses. Correspondingly, the catalytic influence describes the impact of an element on the synthesis of a metabolite in metabolomics data analysis. The phenotypic influence incorporates centrality parameters and represents the topological importance of an element in the curated pathways regulating the phenotype. When the context is unknown or not of interest, we use the network influence as a basic topological parameter of element distance. We provide a detailed explanation of the calculation of each score in the method section. The scores are normalized between −1 and 1, where −1 represents a hypothesized strong negative effect, 0 represents no effect, and 1 represents a strong positive effect from one MIM element to another. For a given phenotype p, the set of elements in the MIM with a nonzero phenotype influence score on p are called **regulators of phenotype *p***.

We introduce an additional weighting factor because the downstream effector and upstream regulator analyses of the IPA software are based solely on the direction of interaction and the sign of fold change. By integrating the network topology-based influence scores between the elements and the enriched term, we improve the accuracy of the enrichment by giving higher weights to topologically more relevant elements. On the other hand, the enriched set of elements is automatically generated using each element with an influence score higher than an absolute threshold. This way, we can integrate thousands of elements with a low impact on the computation time. The resulting enrichment score is generated through association and aggregation of both values (two-dimensional, 2DE) and can either be positive (upregulation) or negative (downregulation) (Figure 5A). We then identify the null distribution by DCE permutation to assess the significance of the enrichment (Figure 5B).

**Figure 5:**
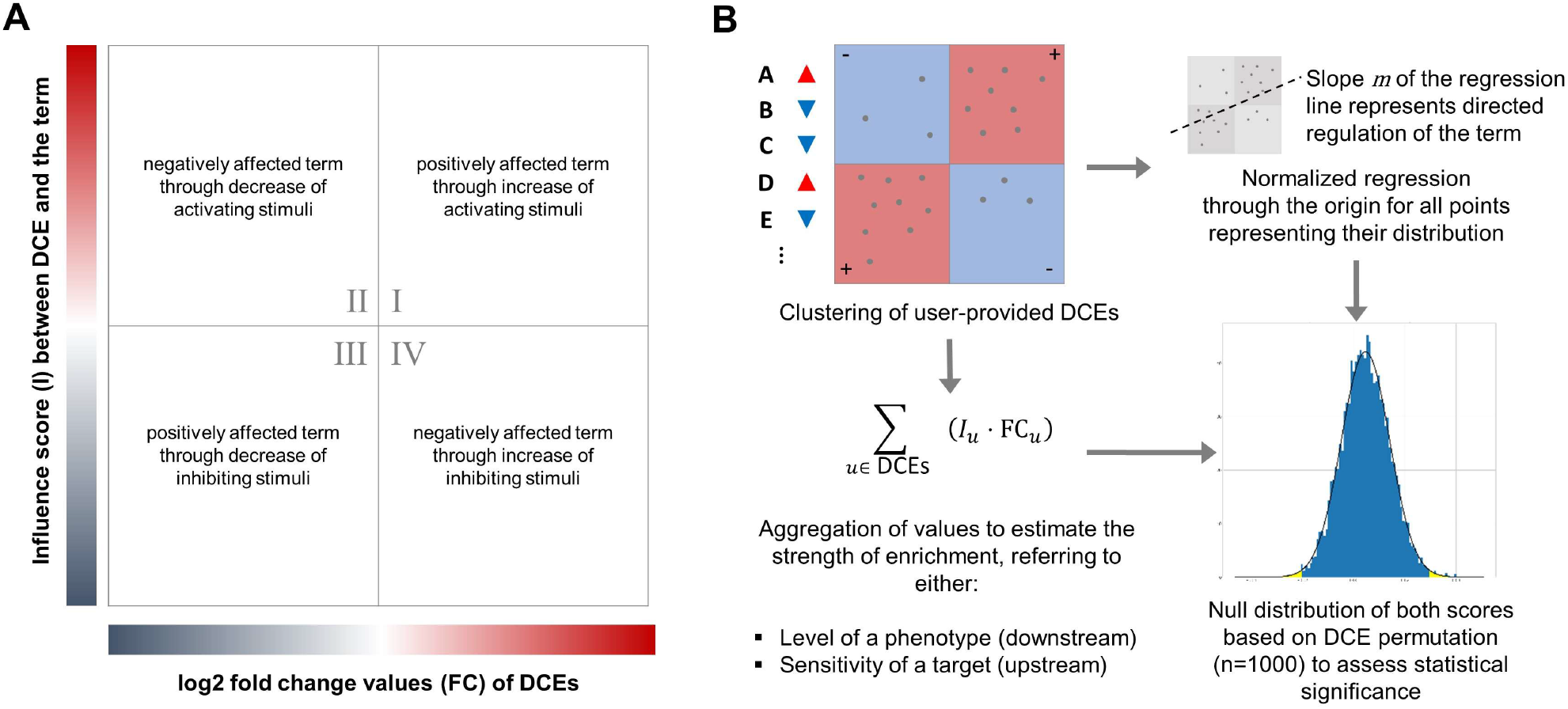
The principle of the two-dimensional enrichment approach (2DE) to identify significantly enriched phenotypes or targets. (A) Log2 fold change values of DCE are compared to their influence score on (downstream) or from (upstream) the term as a weighting factor for the enrichment. (B) Through DCE permutation, we identify the statistical significance of the enrichment.

Some GSEA studies suggest using term label perturbation instead of gene list permutation to avoid scattering the complex co-expression relationships in the data and thus produce more biologically accurate null distributions (Subramanian et al, 2005). However, because the topological relationships in the network define the weighting factors, even a permutation of term labels would not be an entirely realistic distribution. Therefore, we opted for a permutation of DCEs, which is much less computationally expensive because the number of samples in most cases will be less than the number of phenotypes and, more importantly, less than the number of potential targets, which is equal to the total number of elements in the MIM.

#### Downstream Enrichment

Fold changes in DCEs are assumed to be the source or hypothetical cause, and the goal is to identify their effects on other elements in the MIM. This analysis is of particular interest for predicting impacts on **phenotypes**, the enriched terms in this context. Thus, the weighting factors are the influence scores of the DCEs on the phenotype. By aggregating the fold-changes and the influence score values, we obtain a rough estimate of the change in phenotype levels across samples. Because the phenotype level is not an empirical measure, its value is not comparable with other phenotypes. Nevertheless, it provides clues about how the biological process or clinical trait may be regulated across samples. In addition, because the phenotype level is based on DCE aggregation, its value depends on the number of elements considered for the analysis. Therefore, we normalize each phenotype by dividing it by its absolute maximum level across all samples.

#### Upstream Enrichment

The fold changes in DCEs are assumed to be a consequence or “output”. The goal is to identify other elements in the MIM that could be causes and act as enriched terms. We refer to these identified elements as **targets** because they are likely to trigger the observed changes between samples and thus could be the primary driver of disease pathologies. In contrast to downstream enrichment, the weighting factors are the influence scores from targets to DCEs. The definition of target depends on the context and nature of the data but is generally not limited to a specific molecule type. For example, targets can refer to elements associated with changes in the expression profiles of genes in a transcriptomics experiment, changes in the concentrations of metabolites in a metabolomics experiment, or changes in the levels of phenotypes. In addition, targets can either be positive, affecting DCEs according to their FC values, or negative, having the exact opposite effect. Both may be of interest to the user, as suppression of positive targets or activation of negative targets, and vice versa, serves the same purpose. We rank targets according to their **sensitivity** (= true positive rate, i.e., ability to affect DCEs) and **specificity** (= true negative rate, i.e., ability not to affect non-DCEs). Sensitivity is greater than zero for positive targets and less than zero for negative targets. For example, a predicted target with a sensitivity of 1 in a transcriptomics experiment refers to a transcription factor that directly induces the expression of all DEGs with a positive FC value and represses the expression of all DEGs with a negative FC value. In addition, this approach can be adapted to analyze the effect of potential target combinations. By shuffling a set of highest ranked individual targets for all combinations of positive and negative, we again calculate sensitivity and specificity after accumulating their influence scores.

### Enabling data-independent knowledge inference from disease maps

The **Xplore plugin** provides data-independent solutions to answer questions about disease mechanisms *in silico*. It allows users to see changes in downstream phenotypes based on perturbed elements or identify common upstream regulators by defining a desired directed state of phenotypes. Easy-to-use UI elements and color-based visualization facilitate the use of the tools and the interpretability of their results. Because the user inputs involve few elements and serve the purpose of exploration rather than full-fledged analyses, we hold back on statistical analyses and reduce the details in the Xplore plugin to avoid overcomplication. Box 2 highlights the functionalities of the Xplore plugin for disease map exploration and data-independent knowledge inference. The plugin extends the main purpose of disease maps, which is to present knowledge about diseases to the public in an easily understandable form, with tools to perturb molecules or define a biological state and then track its effects or causes in the system.

#### Box 2: Main features of the Xplore plugin.

**Exploring the MIM:** User can access information from the MIM beyond the submaps by selecting an element on the map or manually entering the name. Tables are displayed that give an overview of all direct regulators, targeting elements, and interaction path distances to phenotypes.

**Perturbing elements:** In addition to user defined DCEs, elements that are knocked out (KO) can be specified. These elements are considered removed from the network, resulting in a recalculation of shortest paths and influence scores.

**Highlighting interaction paths:** For each element in the submaps, the shortest path associated with a phenotype can be visualized. If knocked out elements were specified, the path is automatically redirected. Figure S1 shows such a case by disrupting the path of TNF to the phenotype “apoptotic process” through the knock-out of CASP3.

**Visualizing phenotype regulators:** Selecting a phenotype on the map while the plugin is active, automatically highlights all regulators with a colored overlay of red for positive and blue for negative influence scores. Figure S2 shows the submap “Biosynthesis of PIMs and SPMs from AA” after selecting the phenotype “Prostaglandin synthesis”. It shows how hub elements (PGH2) or key enzymes (PTGS2) have the highest scores among all regulators. Additionally, enzymes that catalyze metabolites in the pathway have negative scores due to the consumption of intermediates.

**Inferring phenotypes:** Users set a custom log2 FC value for elements in the MIM through a slider bar (Figure S3), defining them as DCEs. Changing the values of the DCEs automatically updates the predicted level for all phenotypes and visualizes the results as colored overlays on the map.

**Inferring targets:** Users select phenotypes as DCEs by defining a log2 FC value between −1 and 1 representing changes in their activity. Predicted targets, positive and negative, are visualized in a scatter plot ranked by their sensitivity and specificity and can be filtered by their molecule type.

### Inferring downstream phenotypes from a murine colitis model

We developed the **Omics plugin** to enable sophisticated enrichment tools that provide insights into the biological and molecular environment of user-supplied molecular data. Unlike the Xplore plugin, we provide detailed information on the results by graphically displaying the DCEs in each enriched set, intuitively highlighting elements of interest on the submaps, and allowing multiple options for statistical analysis. Users can adjust the parameters of the algorithms or define thresholds for DCEs that fit their data. In addition, we provide an automated optimization function to identify settings with as many filtered DCEs for the highest thresholds as possible. However, users should always interpret the results in the context of the settings by understanding their impact on the calculations. A detailed explanation of the algorithms and their parameters is available in the method section.

To demonstrate data-dependent inference of phenotypes from the plugins, we analyzed a bulk tissue RNA-seq dataset from a murine colitis model and assessed the immune response (Czarnewski *et al*, 2019) (Figure 6). Differentially expressed genes *p*-value < 0.05 were identified for eight time points using DeSeq2 compared to the day-zero control. As input to the Omics plugin, we summarized the results in a tab-delimited .txt file containing the official gene symbol with the respective *log2* fold changes and adjusted *p*-values. For each sample, we identified significantly regulated phenotypes (*p*-value < 0.05). Figure 6C summarizes the results in a heat map showing significant upregulation of cellular inflammatory and lipid mediator related processes between day 6 and day 10. Our results are congruent with the findings of Czarnewski and colleagues, who predicted increased immune cell invasion and cytokine production between day 6 and day 10 based on gene ontology (GO) enrichment.

**Figure 6:**
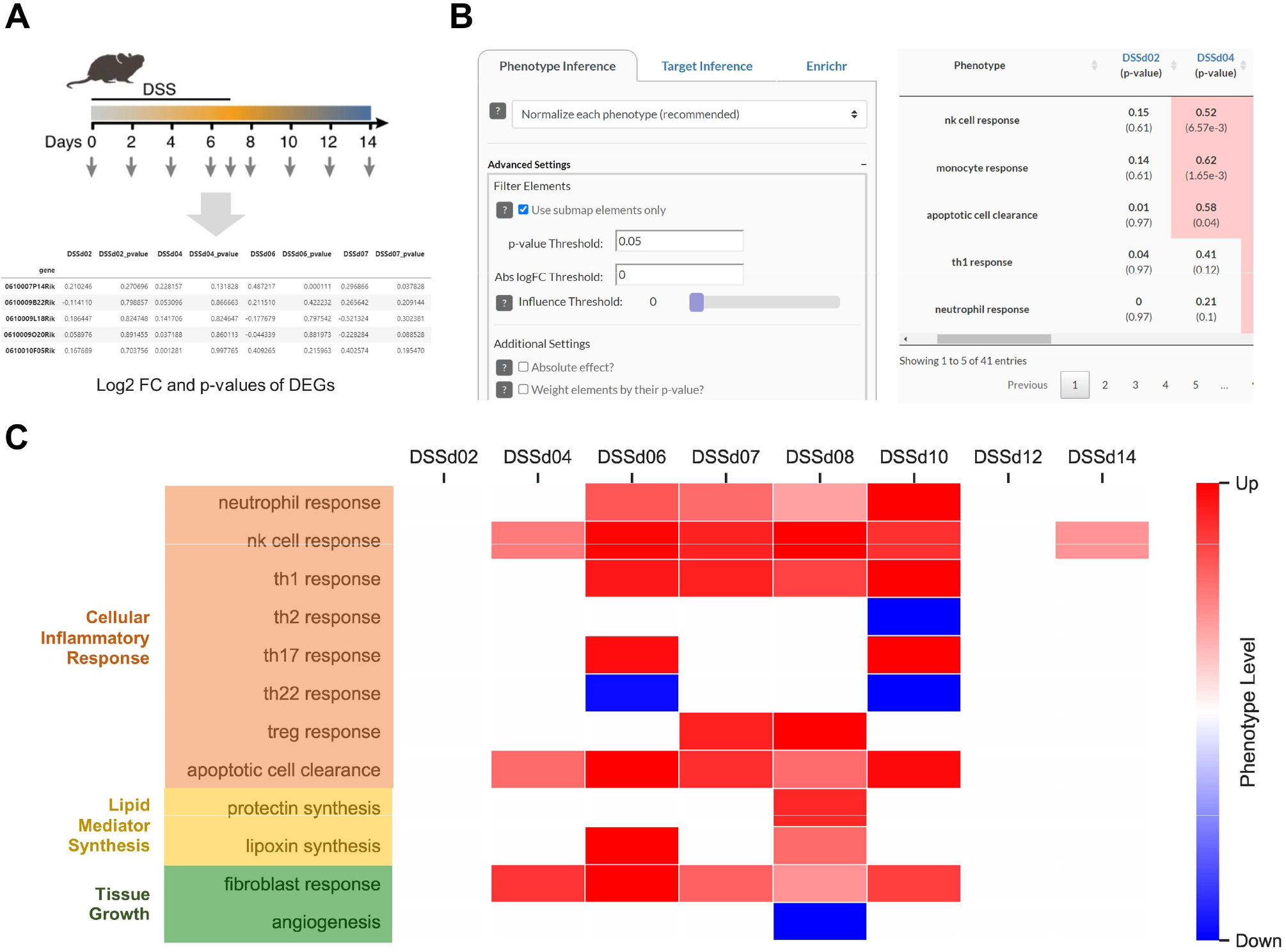
Downstream enrichment analysis with the Omics plugin to identify regulated phenotypes in a murine colitis model. (A) Differential expression was analyzed on colon bulk tissue in mice with DSS-induced colitis for eight different time points (adapted from Czarnewski et al., 2019). (B) In the Omics plugin, we used the phenotype inference by filtering the DEGs for elements from submaps with *p*-value < 0.05. Results are presented in an interactive table, showing predicted levels and *p*-values and creating phenotype regulator plots for each entry. (C) Heatmap of significantly regulated phenotypes in each sample, normalized for each phenotype separately.

### Inferring upstream targets from IFNα-induced differential expression

To demonstrate the target prediction through upstream enrichment, we analyzed RNA-seq data from single-cell B-cells stimulated with IFNα in four different concentrations (1 U, 10 U, 100 U, and 1,000 U) (Mostafavi *et al*, 2016) (Figure 7). Significantly differentially expressed genes *p*-value < 0.05 and *log2* fold change values were loaded in the plugins. We performed an upstream enrichment analysis to identify transcription factor targets with significant interactions to the DEGs in the data. Out of 700 possible TFs in the MIM, we selected the highest scoring TFs, that are also differentially expressed in the supplied dataset. Interestingly, we identified five TFs as significant targets that occurred across all samples, namely IRF7, STAT1, STAT2, EIF2AK2, and PML (Figure 7C). All these targets are known downstream effector targets of IFNα in the literature (Ivashkiv & Donlin, 2014; Gal-Ben-Ari *et al*, 2019; AV *et al*, 2003; R *et al*, 2000). In addition, 5, 20, 48, and 104 DEGs from the data are listed as TFs in the AIR MIM, respectively. All these genes could have been predicted as targets, making a reoccurrence by chance of the top-ranked targets improbable. Figure 7C provides additional insight into the calculation of results by target-regulation plots illustrating the correlation between *log2* fold change values of DEGs and their transcriptional influence scores from STAT1 and IRF7, respectively.

**Figure 7:**
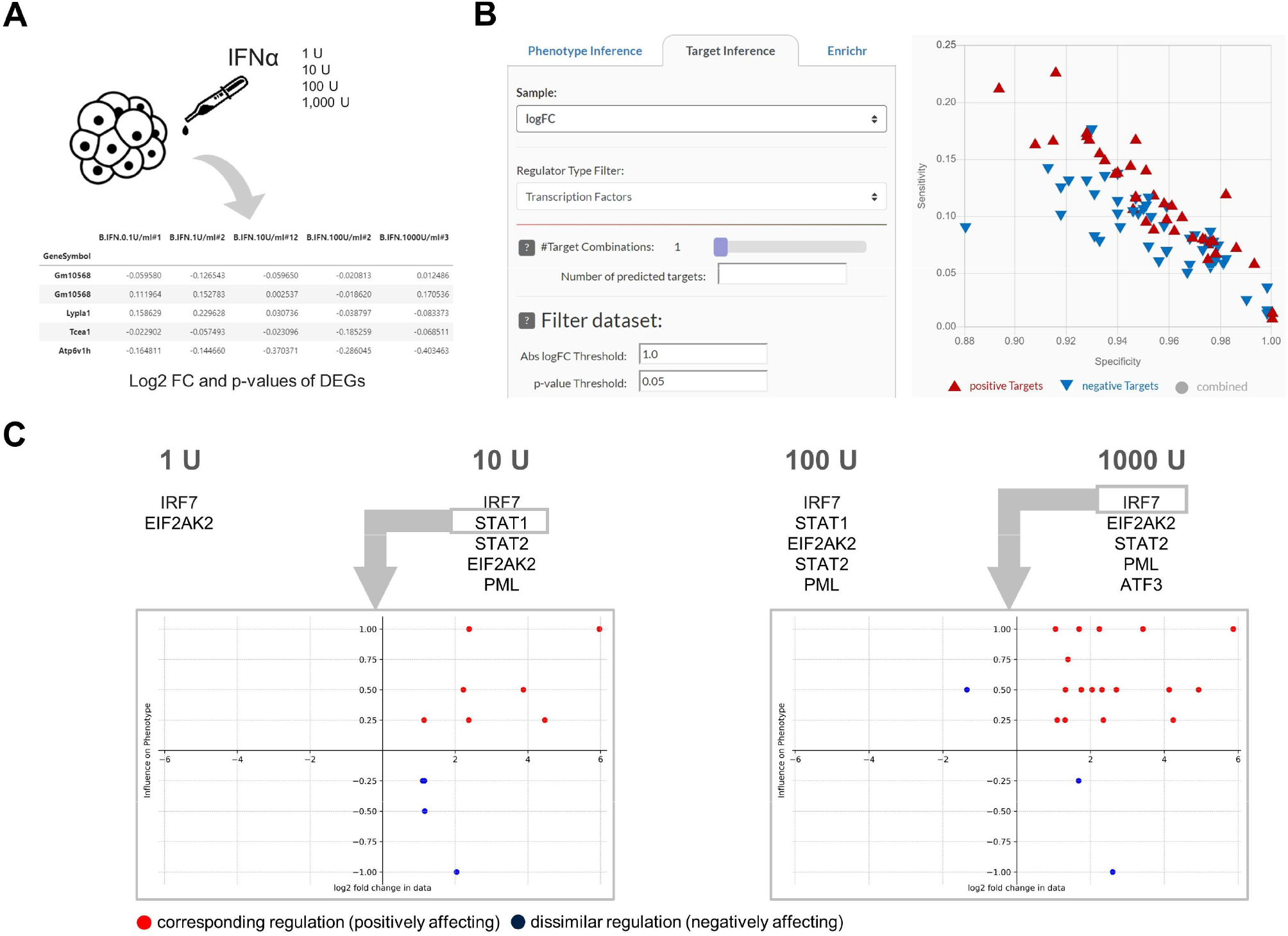
Upstream enrichment analysis with the Omics plugin to identify targets in IFNα induce differential expression. (A) We analyzed the differential expression of IFNα stimulated B-cells for four different concentrations. (B) We used the target inference tool in the Omics plugin by considering only transcript factors and filtering the DEGs for |*log2*| > 1 and *p*-value < 0.05. The plugin presents the results as an interactive scatter plot showing specificity (x-axis) and sensitivity (y-axis) scores. (C) The 5 highest ranked differentially expressed targets (adj. *p*-value < 0.05) for each sample are listed. Additionally, we show two regulations plots for STAT1 and IRF7 each, visualizing *log2* FC values (x-axis) and transcriptional influence scores (y-axis) of regulated DEGs in the data, respectively.

### Comparison with Gene Set Enrichment Analysis (GSEA)

GSEA uses a rank-sum statistic to determine whether an a priori defined group of genes associated with a term is overrepresented at one end of an ordered list of DEGs. This approach has the advantage over simple enrichment statistics such as Fisher’s exact test by including fold changes or weighting factors of DEGs. However, GSEA does not account for the strength or direction of the element’s influence on the phenotype. To determine its differences towards our approach, we ran GSEA on the same case study using phenotypes and their regulators from the AIR as gene sets and the DEGs ranked by their log2 FC value as a gene list input. Figure 8 illustrates which phenotypes are significantly enriched using either of the approaches.

**Figure 8:**
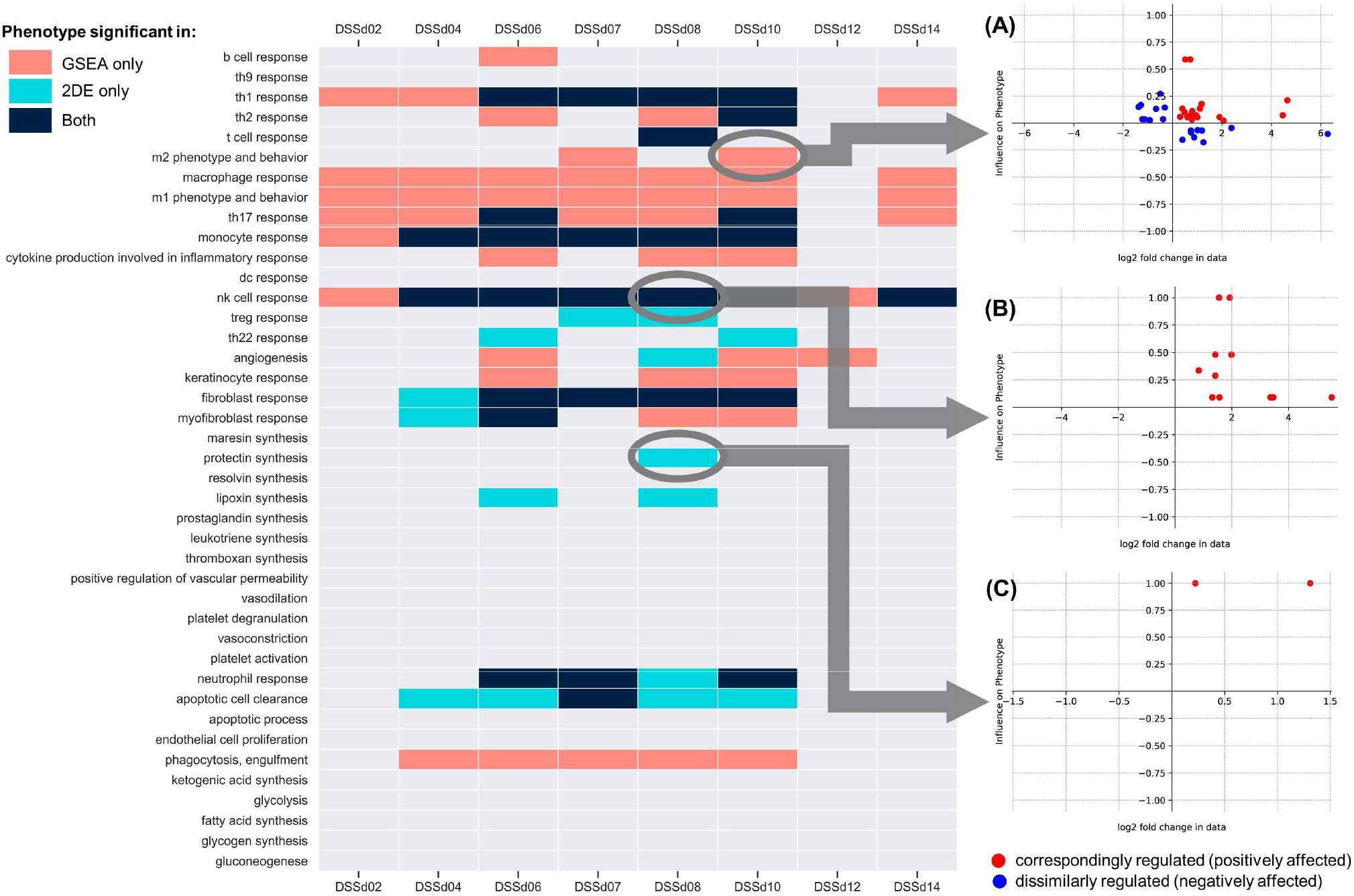
Significantly enriched phenotypes (rows) in each case study sample (columns) predicted by GSEA (red), our 2DE approach (blue) or both (black). (A-C) Phenotype regulator plots show how the two approaches differ in their statistical evaluation for each of the three cases. The running sum statistic on which the GSEA is based predicts phenotypes by distributing phenotypic regulators among the DEGs (x-axis). By integrating the influence score between the regulators and the phenotype (y-axis), our two-dimensional approach excludes phenotypes with ambiguous regulation (A). On the other hand, it detects phenotypes with only a few mapped regulators but with the same directional influence (C).

GSEA identified more phenotypes (68) as significant than our method (37), with many results being overlapping in both approaches (28). We added regulator plots for three cases, i.e., significant in both approaches or only in one of them (Figure 8A-C). GSEA considers a phenotype to be enriched if many DEGs overlap with the phenotype’s regulators. However, these regulators can have opposite effects on the phenotype, making predictions of its aggregated regulation impossible (Figure 8C). In these cases, unlike GSEA, our approach would not predict significance. When both methods are significant, many regulators are clustered in either the phenotype activating (red) or phenotype inhibiting (blue) quadrants (Figure 8B), giving a clear indication of the phenotype’s regulation. In cases where our result was significant but not by GSEA, we see only a few mapped regulators, but they all affect the phenotype similarly (Figure 8B). GSEA does not consider three overlapping DEGs as a statistical significance. However, these three DEGs have a similar influence on the phenotype, demonstrating the advantages of our approach. From a biological point of view, differential expression of a few but essential regulators can strongly influence the phenotype, which would not be detectable by standard enrichment methods.

## DISCUSSION

Disease maps are increasingly valuable knowledgebases for studying disease mechanisms *in silico* and providing researchers and clinicians with an interactive platform for data exploration and visualization. We provide an intuitive solution enabling web-based perturbation experiments and data analysis directly on disease maps.

Our two-dimensional enrichment combines network topology-based relationships between the element and the enriched term, called influence scores, with fold change values from molecular data as weighting factors. The inclusion of both scores allows for much more detailed evaluations by assessing the direction of the response. Even on their own, influence scores are a valuable tool for expanding the information content of disease maps, which provides a visual overview of regulatory processes. Furthermore, influence scores eliminated the need for manual gene set curation by automatically associating elements with existing scores. The two-dimensional approach allows for more accurate predictions of biological regulations compared to other enrichment approaches. In molecular biology, many systems or pathways are regulated by the induction of only a few or even a single key enzyme. Conventional enrichment tools cannot detect these cases where individual changes are distributed among different sets. Our approach does not evaluate the probability that a given element list is overrepresented in the set but whether the accumulated influence of these elements relative to their fold change is statistically significant compared to random permutations. In this way, we can detect enrichments with a small number of associated inputs, allowing more accurate predictions.

By converting large-scale molecular interaction maps from disease maps into enrichment sets of molecule-phenotype or context-specific molecule-molecule associations, we developed a size-independent network-based solution for disease map analysis. We managed to keep computation times to a minimum so that analyses can be performed on the client-side, avoiding the need to upload or store data precluding any data security issues. The approach is highly customizable in that the algorithm for network-based influence score calculations can be adapted for various disease map types without updating the user interface or enrichment part. This customizability improves enrichment capabilities for different data types, e.g., catalytic influence scores for metabolomics data and transcriptional influence scores for transcriptomics data. We successfully addressed many challenges for developing disease map analytic tools, intending to make the methodologies intuitively usable for any interested researcher.

Systems biology approaches should help scientists understand their data and point them to potentially important aspects rather than simply displaying computational results or rankings. The tool we present here focuses on making computations transparent. By incorporating graphical visualization of the DCEs and their weights in the enrichment sets, we provide as much information as possible about the results so that users can make their own interpretations. We plan to extend the plugin suites with features that allow integrating additional data types, such as gene variant or metabolomics data, to enable comprehensive multi-omics analyses on disease maps.

## MATERIAL AND METHODS

### Topological analysis

We developed the plugins using the MIM of the “Atlas of inflammation Resolution”. The AIR MIM is built from 38 manually curated submaps extended with large datasets from public databases describing regulatory interactions of miRNAs, lncRNAs, and transcription factors. We converted the SBML-curated submaps as well as the datasets into a directed graph *G* of a set of elements (vertices *V*(*G*)) and connecting interactions (edges *E*(*G*)). The edges encode whether two elements are linked by (de)activation, up- or downregulation and are defined as a collection of triples *E* ⊂ (*s* × *r* × *t*) consisting of a source element *s* ∈ *V*, a relation *r* ∈ {-1,1}, and a target element *t* ∈ *V*. The full MIM contains more than 6,500 elements connected by a total of over 22,000 interactions. Of the latter, approximately 12,00 are positive, 9,800 are negative. The elements include more than 90 phenotypes, 250 metabolites 4700 proteins, 290 complexes, 460 miRNAs, and 410 lncRNAs. The curation of the AIR and its MIM has been described previously (Serhan *et al*, 2020).

A path *P* in the MIM of the length 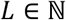 can be written as the sequence 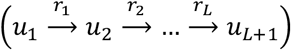 with (*u_i_, r_i_, u_i+1_*) ∈ *E*. The type *T* ∈ {-1,1} of any *P* is defined as (*r*_1_ · *r*_2_ ·… · *r_L_*). The shortest path *SP* between two elements (*u, v*) ∈ *V* is defined as an existing path *P_u,v_* between *u* and *v* where *L*(*P_u,v_*) is minimized. *SP_u,v_* is considered *consistent* if there is no alternative *P_u,v_* with the same length but opposite type. Shortest paths between all pairs of elements were calculated using the Floyd-Warshall algorithm (Floyd, 1962). This algorithm has a complexity of *O*(*V*^3^), which is slower than the modern Dijkstra algorithm with *O*(*E* · *V* + *V*^2^ · *log*(*V*)) when applying on the directed and weighted AIR MIM of *E* ≈ 22,000 and *V* ≈ 6,000. The Floyd-Warshall algorithm was chosen for its ease of implementation and adaptability for the precalculation of values stored in MINERVA. The algorithm calculated *L*(*SP*), *T*(*SP*) for all (*u,v*) ∈ *G* and, for all (*u_s_, v_s_*) originating from submaps, all elements along *P_u_s_,v_s__*. For run-time identification of interaction paths in the plugins, we implemented a simple breadth-first search algorithm.

### Calculation of Influence Scores

The calculation of an influence *I* between two elements *u,v* ∈ *G* in the MIM is based on their connecting paths *P_u,v_*. The routing of the path, however, depends on the context of the analysis. For example, analyzing transcription data, the shortest path leads through transcription factors of *v*. Or, analyzing metabolomics data, the path goes through enzymes in synthesis pathways of *v*. We differentiate between four different types of influence scores through context-specific paths between *u* and *v*. The AIR MIM is a strongly connected graph in which 15% of all pairs of elements are connected by a path. There is a high probability that a random element *u* in the network is connected to, for example, many transcription factors of a given gene *v*. From a biological point of view, an accumulation of all connections between *u* and *v* is closer to actual processes than selecting a particular path based on network parameters. However, due to the interconnectivity of the graph and the decreasing reliability of biological interactions between two elements in molecular networks with increasing distance, it is necessary to define a threshold for *L*(*SP_u,v_*) to be considered as a regulatory interaction. All shortest paths in the MIM have an average length of 6.12 with a standard deviation of 2.3. The probability of two randomly selected elements *u,v* ∈ *V* to be connected by a (*SP_u,v_*) with *L*(*SP_u,v_*) ≤ 2, is 0.2 % and 1.25 % for *L*(*SP_u,v_*) ≤ 3. We decided to use a *L*(*SP_u,v_*) ≤ 2 as a cutoff for topological analyses in the influence scores calculations.

#### Network Influence

(*I_N_*) is based on the distance between *u* and *v* in the MIM (Equation 1):

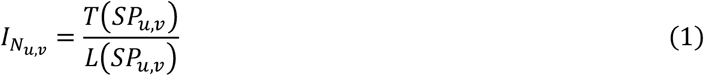

#### Transcriptional influence

(*I_T_*) of *u* on a gene *v* is based on the minimal distance of *u* to *v*’s transcription factors (TF_*v*_) in the MIM (Equation 2). If *u* ∈ TF_*v*_, its influence is equal to the type of interaction between *u* and *v*, i.e., 1 for gene induction or −1 for gene suppression. If *u* ∉ TF_*v*_, its influence is calculated by aggregating the transcriptional influence of each *k* ∈ TF_*v*_ on *v* multiplied by the network influence of *u* on *k*, with *L*(*SP_u,k_*) extended by 1 to ensure *I_T_u,v__* is always smaller than *I_T_k,v__*.

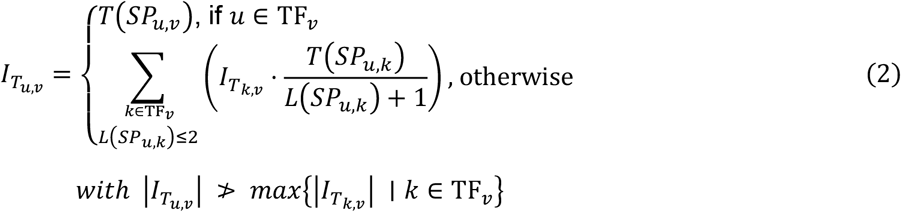

#### Catalytic influence

(*I_C_*) of *u* on a metabolite *v* is based on the minimal distance of *u* to *v*’s synthesizing enzymes (*E_v_*) in the MIM (Equation 3). *E_v_* also includes upstream catalytic enzymes and enzymes that consume *v*. If *u* ∈ *E_v_*, its influence is equal to the type of interaction between *u* and *v*, i.e., 1 for synthesis or −1 for consumption. If *u* ∉ *E_v_*, its influence is calculated by aggregating the catalytic influence of each *k* ∈ *E_v_* on *v* multiplied by the network influence of *u* on *k*, with *L*(*SP_u,k_*) extended by 1 to ensure *I_C_u,v__* is always smaller than *I_C_k,v__*.

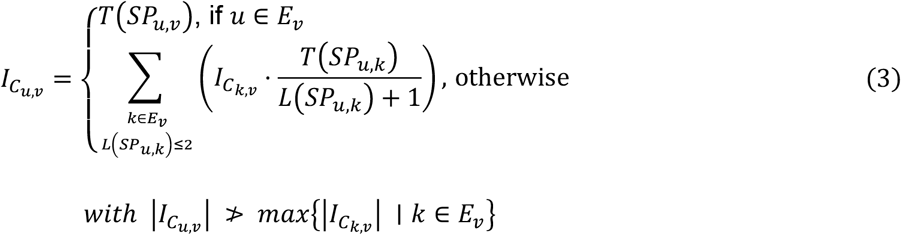

#### Phenotype influence

(*I_P_*) of *u* on a phenotype *v* is based on the topological inclusion of *u* in paths to *v* (Equation 4). *V_s_* ⊂ *V* is the set of elements originating from submaps that contain *v*. If *u* ∈ *V_s_* then its influence is calculated based on the percentage of elements and paths connected with *u. N_P_* is the number of all paths to *v* and *N_P_u__* ⊂ *N_P_* are paths that go through *u. N_G_* is the number of elements connected to *v* and *N_G_u__* ⊂ *N_G_* are elements on the path from *u* to *v*. If *u* ∉ *V_s_*, its influence is calculated by aggregating the phenotype influence of each *k* ∈ *V_s_* on *v* multiplied by the network influence of *u* on *k*, with *L*(*SP_u,k_*) extended by 1.

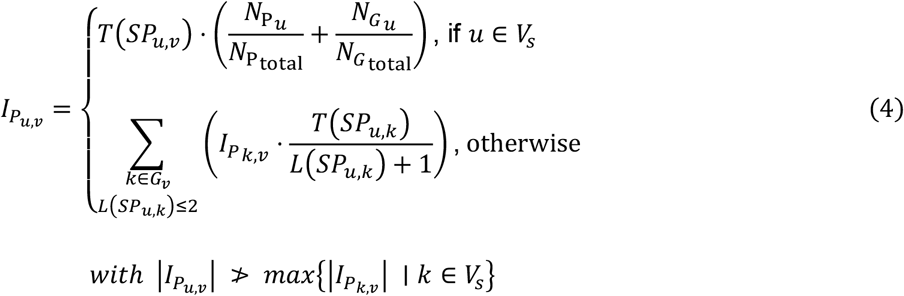

All phenotype influence values are normalized by dividing with the maximum absolute value.

### Aggregating phenotype levels

For each phenotype *v* we calculated the estimated change in activity (= **level**) by aggregating the phenotype influence scores of all regulating elements and their FC value in the given sample (Equation 5).

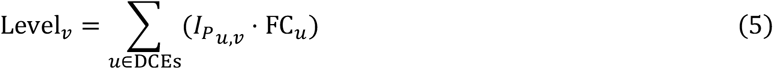

Additionally, we provide information on the **saturation** of the phenotype in the sample, calculated as the percentage of regulators that are DCEs, weighted by their influence score (Equation 6).

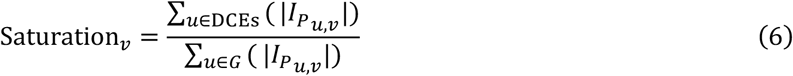

### Ranking of targets

Targets for a set of DCEs are ranked by their **sensitivity** and **specificity**, calculated as follows for a specific target *V*. Sensitivity (= true positive rate, Equation 7) will be 1 (= positive target) if the influence of *v* on every DCE is 1. Conversely, the sensitivity will be −1 (= negative target) if the influence of *v* on every DCE is −1. Specificity (= true negative rate, Equation 8) will be 1 if the influence of *v* on every non-DCE is 0.

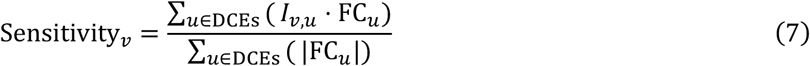

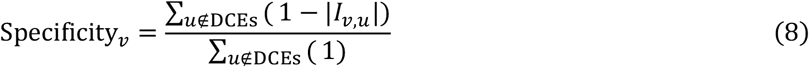

### Statistical evaluation

For each sample in the supplied data, *n* random sets (*n* = 1000 by default) of DCEs (DCE_1_, DCE_2_,…, DCE_*n*_) are generated with an equal number and values of the filtered significant *log2* FC values as the original DCEs, randomized among all MIM elements of the same type (e.g., genes or metabolites). Depending on the context and type of analysis, a score is calculated for each random set and its distribution identified through Gaussian fitting. Then, using the mean μ and standard deviation σ of the distribution, a *p*-value is calculated as the probability to achieve the same or higher score than the original DCE set at random (one- or two-sided) to determine its significance. Adjustment for multiple testing is then performed using false discovery rate (FDR)-correction by *Benjamini–Hochberg* to generate the adjusted *p*-value (Benjamini & Hochberg, 1995).

The *Sensitivity* value is used as the score for the target prediction (forward tracing) statistics, while for the phenotype prediction (backward tracing), we provide three different scores:

1. **Level-based**: The score is defined as the level of the phenotype, providing a statistical evaluation of the regulation strength.
2. **Enrichment-based**: The score is defined as the absolute level of the phenotype using absolute FC and Influence values.

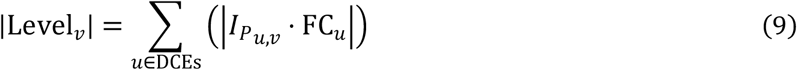
3. **Distribution-based**: A linear regression through the origin is performed for all points (x,y) in

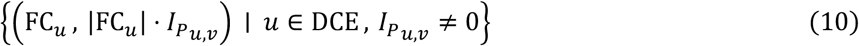 The score is defined as the normalized slope of the regression line *m* (*m*_norm_) between −1 (45° decline) and 1 (45° incline):

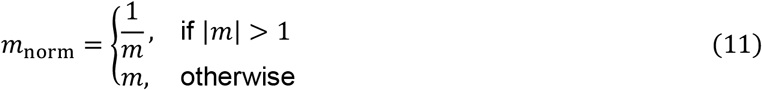 Using the distribution as a statistical parameter, the significance is not based on the number of DCEs but rather their influence on the phenotype.

### Processing RNA-seq data

For the murine colitis model, Kallisto raw counts were fetched from GEO (Accession number GSE131032). The data were analyzed using the R Deseq2 package comparing each sample with the day zero control. For IFNα stimulated B-cells, we used GEO’s GEO2R to directly compute *log2* FC values and adjusted *p*-value for all samples vs the control (Accession number GSE75194). For each dataset, its results were summarized in a tab-separated text file containing the gene name in the first column together with the *log2* fold change and adjusted *p-*value for each comparison, respectively, as additional columns.

### Reagents and Tools Table

| Resource | Reference or Source | Identifier or Catalog Number |
| --- | --- | --- |
| **Software**<br>*Include version where applicable* |  |  |
| npm v6.14.4 | https://www.npmjs.com/ |  |
| RStudio v1.4.1106 | http://www.rstudio.com/ |  |
| DeSeq2 for R v1.28.1 | (Love *et al*, 2014) |  |
| CellDesigner v4.4.2 | (Funahashi *et al*, 2007) |  |
| GSEAPy v0.10.5 | https://gseapy.readthedocs.io/ |  |

## Acknowledgments

MINERVA is developed by the Luxembourg Centre for Systems Biomedicine (LCSB). We are grateful for the support provided by Piotr Gawron and Marek Ostaszeweski in the development and deployment of the plugins.

## Author contributions

M. Hoch developed the tools and performed the analyses. All authors contributed to the scientific content, helped writing the text, and approved the submitted version. Furthermore, all authors contributed to the interpretation and quality control of the information contained in the AIR.

## References

Aghamiri SS, Singh V, Naldi A, Helikar T, Soliman S & Niarakis A (2020) Automated inference of Boolean models from molecular interaction maps using CaSQ. Bioinformatics 36: 4473–4482

AV C, P L, PP P & B R (2003) Promyelocytic leukemia protein mediates interferon-based antiherpes simplex virus 1 effects. J Virol 77: 7101–7105

Benjamini Y & Hochberg Y (1995) Controlling the False Discovery Rate: A Practical and Powerful Approach to Multiple Testing. J R Stat Soc Ser B 57: 289–300

Bindea G, Mlecnik B, Hackl H, Charoentong P, Tosolini M, Kirilovsky A, Fridman W-H, Pagès F, Trajanoski Z & Galon J (2009) ClueGO: a Cytoscape plug-in to decipher functionally grouped gene ontology and pathway annotation networks. Bioinformatics 25: 1091–1093

Chen EY, Tan CM, Kou Y, Duan Q, Wang Z, Meirelles G, Clark NR & Ma’ayan A (2013) Enrichr: interactive and collaborative HTML5 gene list enrichment analysis tool. BMC Bioinformatics 14: 128

Czarnewski P, Parigi SM, Sorini C, Diaz OE, Das S, Gagliani N & Villablanca EJ (2019) Conserved transcriptomic profile between mouse and human colitis allows unsupervised patient stratification. Nat Commun 10: 2892

Floyd RW (1962) Algorithm 97: Shortest path. Commun ACM 5: 345

Fujita KA, Ostaszewski M, Matsuoka Y, Ghosh S, Glaab E, Trefois C, Crespo I, Perumal TM, Jurkowski W, Antony PMA, et al (2014) Integrating Pathways of Parkinson’s Disease in a Molecular Interaction Map. Mol Neurobiol 49: 88–102

Funahashi A, Morohashi M, Matsuoka Y, Jouraku A & Kitano H (2007) CellDesigner: A Graphical Biological Network Editor and Workbench Interfacing Simulator. In Introduction to Systems Biology pp 422–434. Totowa, NJ: Humana Press

Gal-Ben-Ari S, Barrera I, Ehrlich M & Rosenblum K (2019) PKR: A Kinase to Remember. Front Mol Neurosci 0: 480

Gawron P, Ostaszewski M, Satagopam V, Gebel S, Mazein A, Kuzma M, Zorzan S, McGee F, Otjacques B, Balling R, et al (2016) MINERVA—a platform for visualization and curation of molecular interaction networks. npj Syst Biol Appl 2: 16020

Helikar T, Kowal B, McClenathan S, Bruckner M, Rowley T, Madrahimov A, Wicks B, Shrestha M, Limbu K & Rogers JA (2012) The Cell Collective: Toward an open and collaborative approach to systems biology. BMC Syst Biol 6: 96

Hong G, Zhang W, Li H, Shen X & Guo Z (2014) Separate enrichment analysis of pathways for up- and downregulated genes. J R Soc Interface 11: 20130950

Ivashkiv LB & Donlin LT (2014) Regulation of type I interferon responses. Nat Rev Immunol 14: 36–49

Jiao X, Sherman BT, Huang DW, Stephens R, Baseler MW, Lane HC & Lempicki RA (2012) DAVID-WS: A stateful web service to facilitate gene/protein list analysis. Bioinformatics 28: 1805–1806

Kanehisa M (2000) KEGG: Kyoto Encyclopedia of Genes and Genomes. Nucleic Acids Res 28: 27–30

Keating SM, Waltemath D, König M, Zhang F, Dräger A, Chaouiya C, Bergmann FT, Finney A, Gillespie CS, Helikar T, et al (2020) <scp>SBML</scp> Level 3: an extensible format for the exchange and reuse of biological models. Mol Syst Biol 16: e9110

Kitano H, Funahashi A, Matsuoka Y & Oda K (2005) Using process diagrams for the graphical representation of biological networks. Nat Biotechnol 23: 961–966

Krämer A, Green J, Pollard J & Tugendreich S (2014) Causal analysis approaches in Ingenuity Pathway Analysis. Bioinformatics 30: 523–530

Love MI, Huber W & Anders S (2014) Moderated estimation of fold change and dispersion for RNA-seq data with DESeq2. Genome Biol 15: 550

Martens M, Ammar A, Riutta A, Waagmeester A, Slenter DN, Hanspers K, A. Miller R, Digles D, Lopes EN, Ehrhart F, et al (2021) WikiPathways: connecting communities. Nucleic Acids Res 49: D613–D621

Mazein A, Knowles RG, Adcock I, Chung KF, Wheelock CE, Maitland-van der Zee AH, Sterk PJ & Auffray C (2018a) AsthmaMap: An expert-driven computational representation of disease mechanisms. Clin Exp Allergy 48: 916–918

Mazein A, Ostaszewski M, Kuperstein I, Watterson S, Le Novère N, Lefaudeux D, De Meulder B, Pellet J, Balaur I, Saqi M, et al (2018b) Systems medicine disease maps: community-driven comprehensive representation of disease mechanisms. npj Syst Biol Appl 4: 21

Mostafavi S, Yoshida H, Moodley D, LeBoité H, Rothamel K, Raj T, Ye CJ, Chevrier N, Zhang S-Y, Feng T, et al (2016) Parsing the Interferon Transcriptional Network and Its Disease Associations. Cell 164: 564–78

Ostaszewski M, Gebel S, Kuperstein I, Mazein A, Zinovyev A, Dogrusoz U, Hasenauer J, Fleming RMTT, Le Novère N, Gawron P, et al (2019) Community-driven roadmap for integrated disease maps. Brief Bioinform 20: 659–670

Parton A, McGilligan V, Chemaly M, O’Kane M & Watterson S (2019) New models of atherosclerosis and multi-drug therapeutic interventions. Bioinformatics 35: 2449–2457

R L, WC A, WS Y, N H & PM P (2000) Regulation of the promoter activity of interferon regulatory factor-7 gene. Activation by interferon snd silencing by hypermethylation. J Biol Chem 275: 31805–31812

Serhan CN, Gupta SK, Perretti M, Godson C, Brennan E, Li Y, Soehnlein O, Shimizu T, Werz O, Chiurchiù V, et al (2020) The Atlas of Inflammation Resolution (AIR). Mol Aspects Med 74: 100894

Singh V, Ostaszewski M, Kalliolias GD, Chiocchia G, Olaso R, Petit-Teixeira E, Helikar T & Niarakis A (2018) Computational Systems Biology Approach for the Study of Rheumatoid Arthritis: From a Molecular Map to a Dynamical Model. Genomics Comput Biol 4: 100050

Subramanian A, Tamayo P, Mootha VK, Mukherjee S, Ebert BL, Gillette MA, Paulovich A, Pomeroy SL, Golub TR, Lander ES, et al (2005) Gene set enrichment analysis: a knowledge-based approach for interpreting genome-wide expression profiles. Proc Natl Acad Sci U S A 102: 15545–50

Warden CD, Kanaya N, Chen S & Yuan Y-C (2013) BD-Func: a streamlined algorithm for predicting activation and inhibition of pathways. PeerJ 1: e159

Zito A, Lualdi M, Granata P, Cocciadiferro D, Novelli A, Alberio T, Casalone R & Fasano M (2021) Gene Set Enrichment Analysis of Interaction Networks Weighted by Node Centrality. Front Genet 12: 577623

