## Supplementary figures and images for "Network- and Enrichment-based Inference of Phenotypes and Targets from large-scale Disease Maps"

### Figure S1

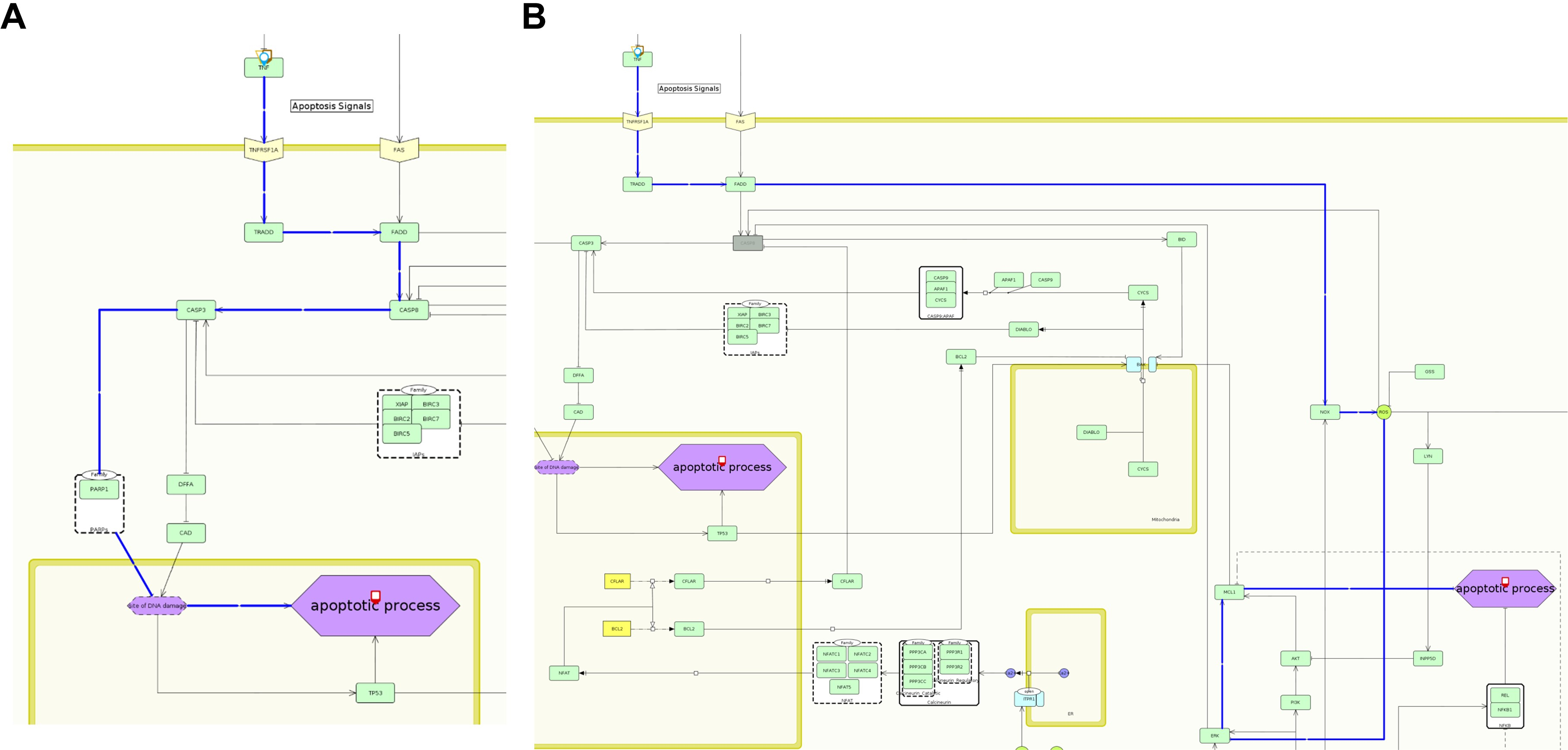

### Figure S2

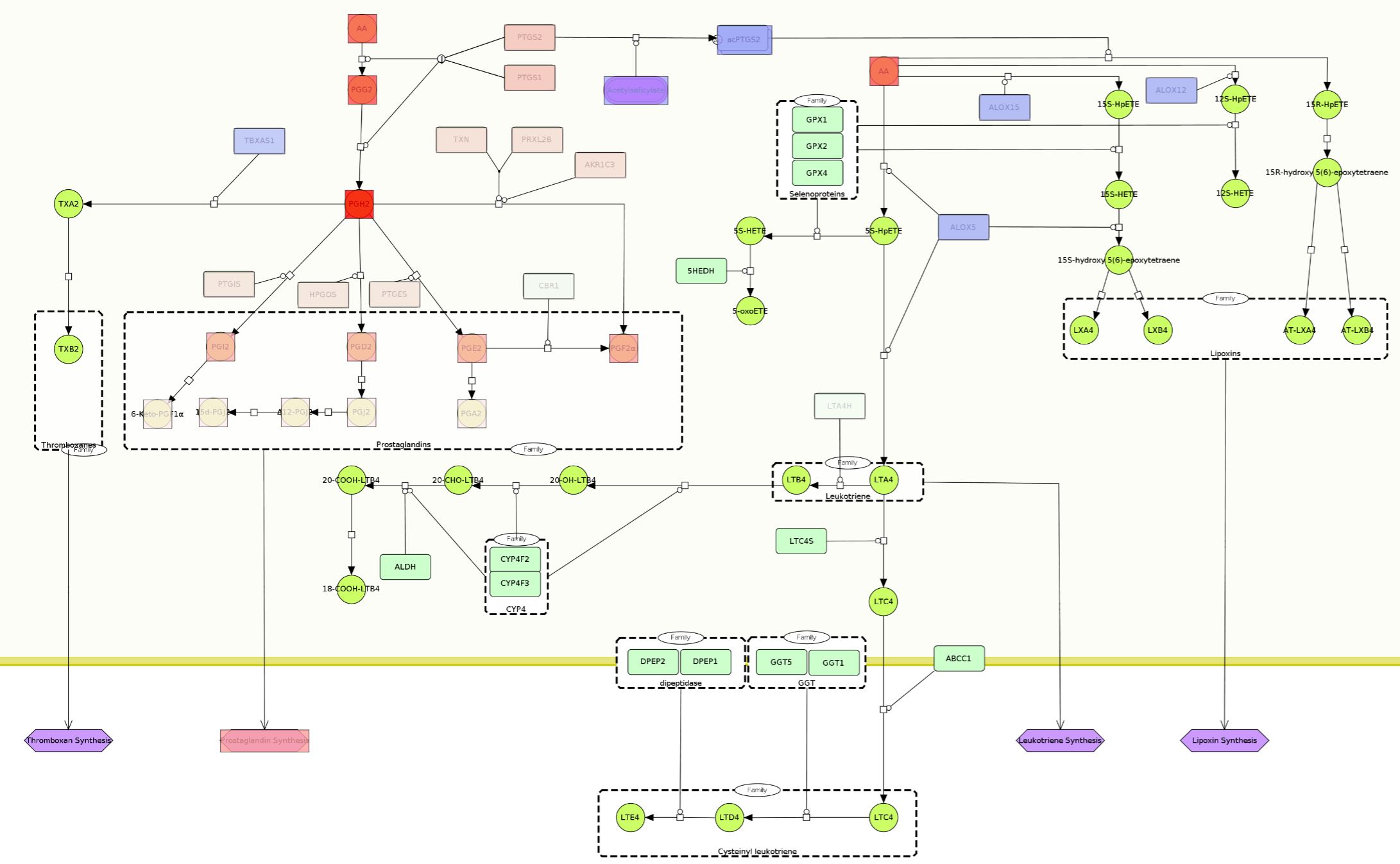

### Figure S3

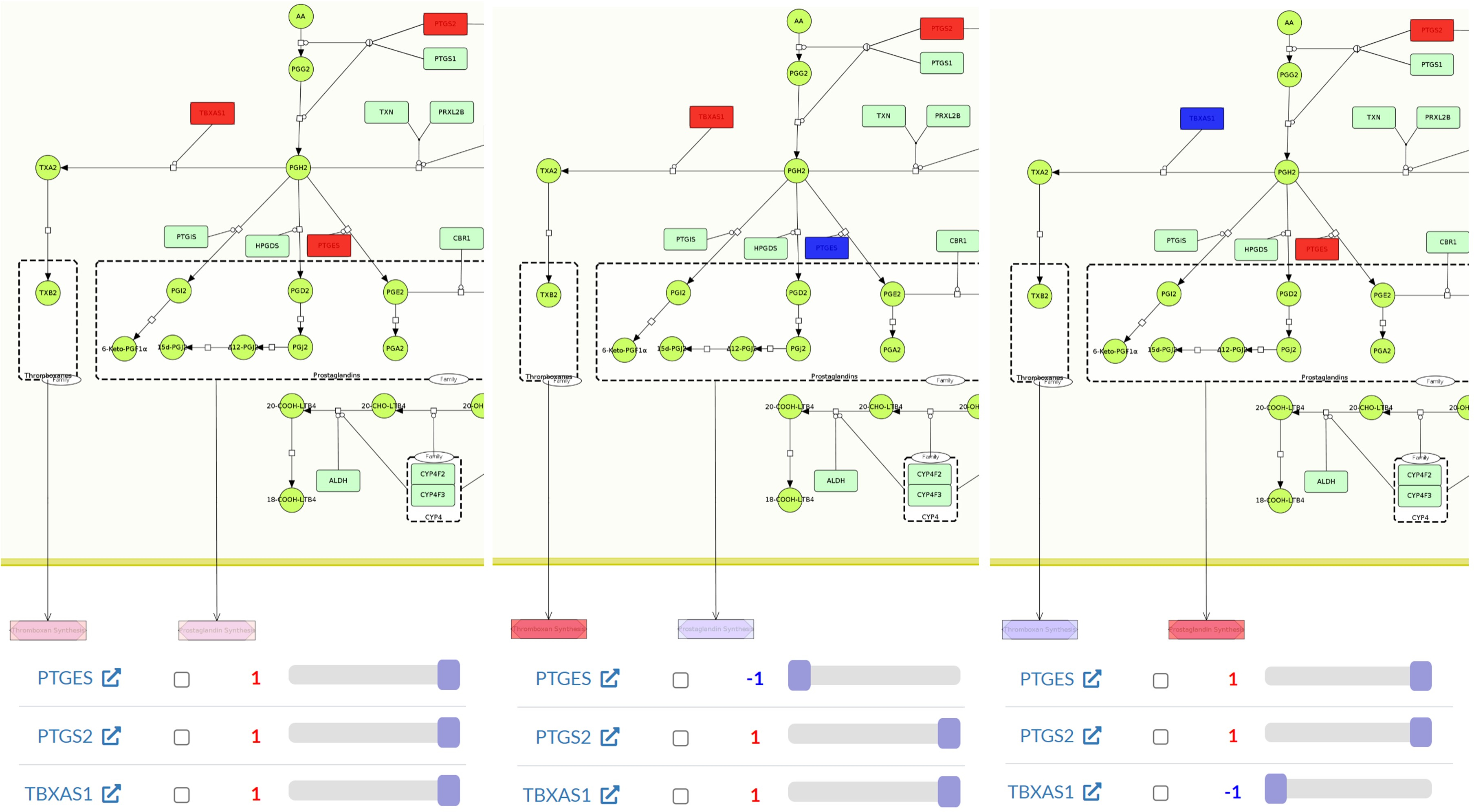
